# Branching architecture constrains the accumulation of somatic mutations in trees

**DOI:** 10.1101/2025.02.21.639457

**Authors:** Frank Johannes

## Abstract

Trees are long-lived plants that develop highly branched shoot systems over time. During growth, somatic mutations accumulate along these branching structures and can become fixed in reproductive tissues such as flowers and fruits. Because mature trees produce tens of thousands of terminal branches, each harboring potentially mutated gametes, limiting the accumulation of somatic mutations is critical to avoid mutational meltdown and inbreeding depression. Although recent evidence suggests that long-lived plants have evolved mechanisms to suppress mutation accumulation, the developmental basis for this remains unclear. Here we develop a theoretical model that connects crown development with cell lineage sampling to show that *branching architecture* can strongly suppress somatic mutation accumulation in trees, often to the same extent as reducing the mutation rate itself. Specifically, tree forms that promote developmental path-sharing among branches suppress the number of unique mutational lineages in the crown. We find that this architectural effect can alter mutation burden by orders of magnitude, even when mutation rates and terminal branch numbers remain constant. These insights suggest that branching strategies may evolve not only to optimize growth and resource allocation, but also to limit the accumulation of somatic variants during ontogeny.

---

Trees are long-lived organisms that gradually develop complex, highly branched shoot systems over time (Fig. 1A). In many species, mature individuals can sustain tens of thousands of terminal branches, each derived from a shoot apical meristem (SAM), a small, self-renewing population of stem cells located at the shoot apex. Because the SAM gives rise to the entire shoot system, somatic mutations arising in this region can be perpetuated across large sectors of the plant body (Chen et al. (2024); Tomimoto and Satake (2023); Iwasa et al. (2023); Johannes (2025)). These mutations may accumulate over time and be retained in both vegetative structures (e.g., leaves, stems, fruit) and reproductive tissues. As a result, they can influence not only the development and physiology of the individual tree, but also its genetic contribution to the next generation (Schoen and Schultz (2019); Quiroz et al. (2023)).

**Figure 1:**
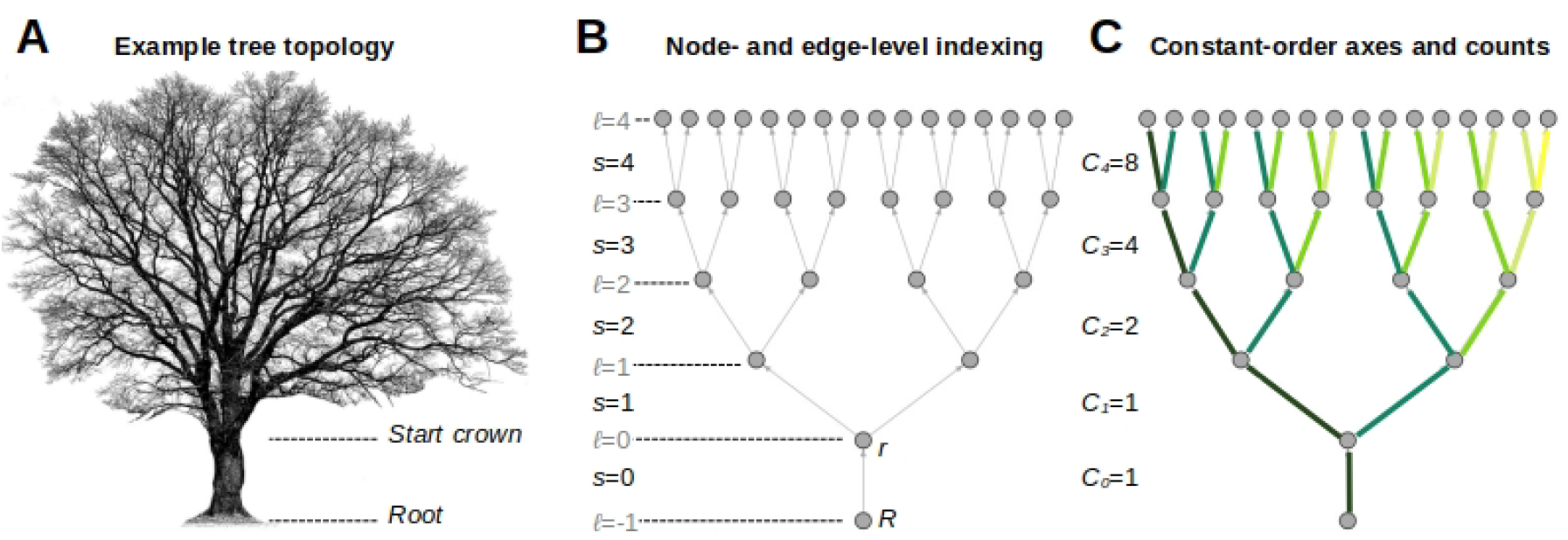
Graph presentation of the tree crown. **A**. Trees develop complex, highly branched shoot systems as they grow. An example of an oak tree is shown. Mature specimen can sustain hundreds of thousands of terminal branches. **B**. The branching structure of a tree can be presented as a connected, acyclic graph with edges representing internode segments and nodes branch points. Node *R* represents the actual root of the tree and node *r* marks the start of the tree crown. The crown itself can be modeled as a perfectly balanced binary tree, where each internal node has two children. The hierarchical structure of the tree can be quantified by its depth, *D* (here *D* = 4). It represents the maximum distance from the root to the terminal nodes. Node levels are indexed as *ℓ* ∈ {− 1, 0, …, *D*} and edge levels as *s* ∈ {0, …, *D*}. **C**. We define constant-order axes as unique paths starting at level *s* and ending in one of the terminal nodes without diverting into lateral branches. The color coding highlights these axes. We use *C*_*s*_ to count the number of unique constant-order axes for each level *s*.

Plants with long life spans appear to have evolved strategies to mitigate the detrimental effects of mutation load and reduce the risk of inbreeding depression. These include low selfing rates and large effective population sizes, both of which help maintain genetic diversity and minimize inbreeding coefficients (Lesaffre (2021); Scofield and Schultz (2006); Schoen and Schultz (2019)). In addition to these population-level adaptations, emerging evidence points to intrinsic mechanisms that actively slow the accumulation of somatic mutations per unit time (Lanfear et al. (2013)). This is apparent in the reduced somatic mutation rate seen in long-lived tree species like *O. fragrans*, which is about two orders of magnitude lower per year than in shorter-lived species like *S. suchowensis*(Johannes (2024); De La Torre et al. (2017)). A similar trend has been observed at the epigenetic level, where epimutation rates, measured as stochastic gains and losses of DNA methylation, are negatively correlated with generation time. This parallel reduction suggests a shared underlying biological mechanism that affects both genetic and epigenetic mutation processes (Johannes (2024)). However, the exact nature of this mechanism remains unknown. One hypothesis is that long-lived trees simply slow meristematic cell divisions per unit time (Lanfear et al. (2013)). This mechanism would reduce errors arising from DNA synthesis and DNA methylation maintenance during replication. Although direct *in vivo* validation of this hypothesis remains challenging, it is strongly supported by the recent observation that somatic (epi)mutation rates correlate positively with growth rates and cell division frequency in vegetative tissues (Lanfear et al. (2013); Zhou et al. (2024)).

The link between growth dynamics and somatic mutation accumulation suggests a broader connection between molecular evolution and life-history traits. Since SAM-derived mutations are primarily incorporated into lateral organs during branch initiation, the spatial and temporal dynamics of branching may present a major rate limiting step for the accumulation of mutations in the tree crown, a possibility that has received little attention. Here, we present a theoretical analysis to investigate this relationship. We demonstrate that branching architectures and branch age distributions can substantially modulate somatic mutation burden, with effects comparable to reducing the mutation rate itself. Tree structures that maximize developmental path sharing, where many terminal branches trace back to a common meristematic lineage, tend to suppress mutational diversity. In some cases, these architectural features can reduce the mutation burden by orders of magnitude, even when baseline mutation rates and terminal branch numbers remain unchanged.

Our findings have major implications for understanding the rate and topological distribution of somatic mutations in trees, and highlight life-history traits as key determinants of intra-organism somatic heterogeneity.

## Theory and discussion

### Modeling the spatio-temporal architecture of a tree crown

#### A graph representation of the crown

We model the tree crown as a rooted, directed acyclic graph *A* = (*V, E*), in which each vertex (*V*) represents a branch point and each directed edge (*E*) an internodal segment (Fig. 1A-B, Table1). A special root node *R* connects the basal segment (*s* = 0) with the first crown branch point *r* at level 0. Above this, we assumed that the graph forms a perfectly balanced binary structure of depth *D*, with level *ℓ* containing 2^*ℓ*^ vertices (*ℓ* = 0, 1, …, *D*), although this assumption can be easily relaxed. In this structure each vertex at level *s* − 1 has exactly two children at level *s*, so that each path from *R* to a terminal meristem traverses precisely *D* + 1 segments indexed by *s* = 0, 1, …, *D*. The number of remaining branch points above segment *s* can be calculated as

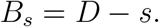

**Table 1:** Key notation and indexing for the tree-crown model.

| Symbol | Meaning |
| --- | --- |
| $D$ | Crown depth (number of bifurcation levels) |
| $\ell$ | Node level, $\ell = -1, 0, \dots, D$ |
| $s$ | Segment index, $s = 0, \dots, D$ |
| $B_s$ | Downstream branch points above $s$, $D - s$ |
| $C_s$ | Constant-order axes through segment $s$ |
| $T$ | Total developmental time per path |
| $\lambda$ | Exponential tilt for time allocation |
| $w_s(\lambda)$ | Proportion of $T$ on segment $s$ |
| $t_s$ | Time on segment $s$, $T w_s(\lambda)$ |

The counts *B*_*s*_ will later become useful when indexing the number of cellular bottlenecks a mutation on segment *s* must clear to reach a terminal meristem.

#### Internode orders and constant-order axes

When modeling shoot branching, it is important to distinguish the main growth axis from lateral shoots. To do this, we assign each internode segment an *order* : the basal segment *s* = 0 has order zero; at each branching point the continuation segment retains its parent’s order, while the lateral segment’s order increments by one. A *constant-order axis* is any maximal chain of continuation segments along which the order remains unchanged until a terminal tip is reached (Fig. 1C, Table1). Denote by *C*_*s*_ the number of constant-order axes that pass through segment *s*. In a perfectly balanced binary tree of depth *D*, one finds

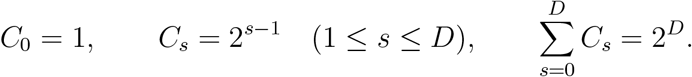

#### Developmental time allocation

Having established the basic tree topology, let us now use *T* to denote the total developmental time from root *R* to tip *D*. It can be measured in terms of years, number of mitotic cell divisions or any other unit of time. We partition *T* across the *D* + 1 internode segments by introducing an exponential tilt function. Define

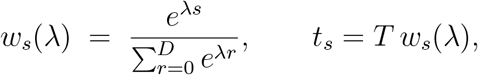

so that ∑_*s*_ *w*_*s*_(*λ*) = 1 and ∑_*s*_ *t*_*s*_ = *T*. Here, *t*_*s*_ is the time assigned to internode segment *s*. By modulating the parameter *λ* this tilt function allows us to explore diverse growth regimes. For example, in the limiting case where *λ* → −∞, we have that

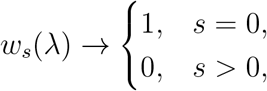

meaning that all time is concentrated on the basal segment. This growth regime produces a “stick-like” tree architecture (Fig.2B, left panel). When *λ* = 0, *w*_*s*_ = 1*/*(*D* + 1) for every *s*, yielding uniform time allocation (Fig.2B, right panel). At the other extreme, *λ* → +∞,

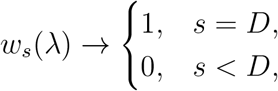

so that all time is allocated to the terminal segment. This growth regime yields a “star-like” tree architecture (Fig.2B, middle panel). Hence, by tuning *λ*, this oneparameter family smoothly shifts developmental strategies from trunk-dominated through uniform to tip-dominated growth.

**Figure 2:**
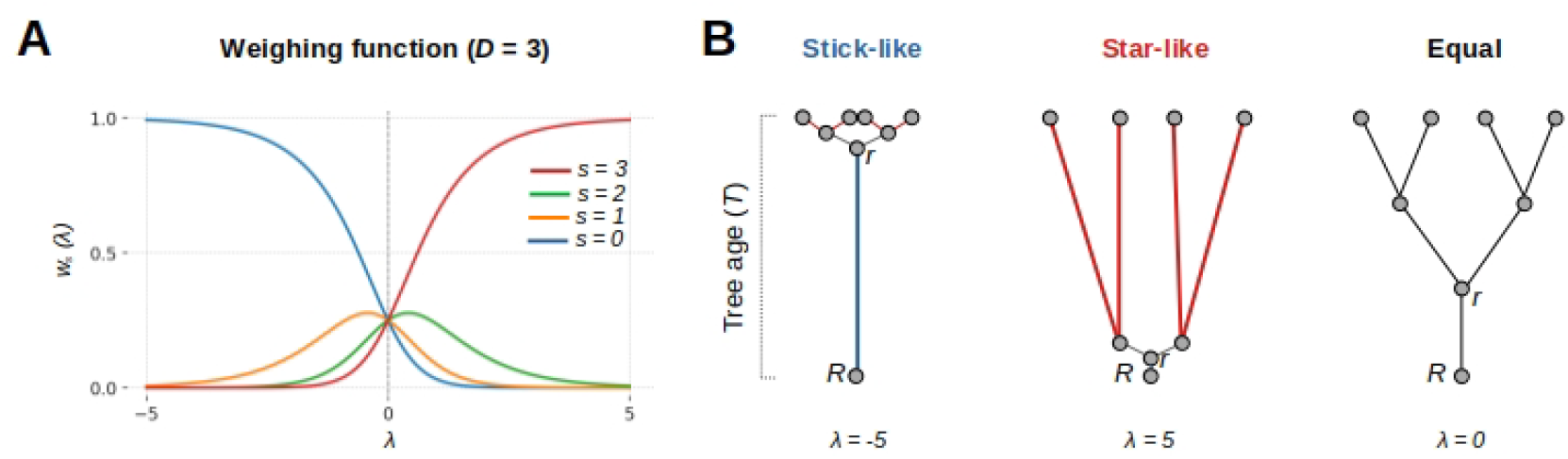
Developmental time allocation. **A**. Exponential tilt function *w*_*s*_(*λ*) allocates time *t*_*s*_ to internode segments *s* (colored lines). For *D* = 3, the plot shows how the parameter *λ* governs time allocation. For large negative *λ*, developmental time is allocated to the root segment *s* = 0 (blue line); for large positive *λ* time is shifted toward the terminal segment *s* = 3 (red line) and for *λ* = 0, time is allocated equally across segments (intersection points of lines). **B**. Topological consequences of the time allocation regimes described in A.

### Mutation sampling and propagation

Our goal is to model how somatic mutations accumulate across the tree architectures described above. Among all possible somatic variants, those arising within the SAM are the most consequential, as all aerial cell lineages originate from this tissue. SAM-derived mutations can spread across large sectors of the plant body, especially when transmitted into axillary meristems (AMs) that establish new lateral branches or organs (Fig.3). These mutations are more likely to reach high cellular frequencies, making them detectable via sequencing, and have greater potential for functional impact and entry into the germline. In contrast, mutations arising outside the SAM are typically confined to small, localized sectors, are challenging to access with genomic tools, and are arguably of less evolutionarily significance.

#### Cellular model of shoot branching

To understand how SAM-derived mutations spread throughout the plant, we must posit a cellular model of shoot branching that captures how SAM lineages contribute to lateral meristem formation. The SAM is a stratified, self-renewing structure composed of two tunica layers (L1, L2) and one corpus layer (L3) (Lyndon (1998)). Each layer is maintained by a small number of apical stem cells (ASCs) (typically *m* = 3–5 per layer in species such as *Arabidopsis*, tomato, or seagrass; Burian et al. (2016); Yu et al. (2024)) arranged at the shoot apex (Fig. 3A). During internode growth, ASCs divide asymmetrically: one daughter retains its stem cell identity, while its sister is displaced into the periphery and contributes to organogenesis. This produces coherent clonal sectors within each layer, each tracing back to a single ASC (Fig. 3B).

**Figure 3:**
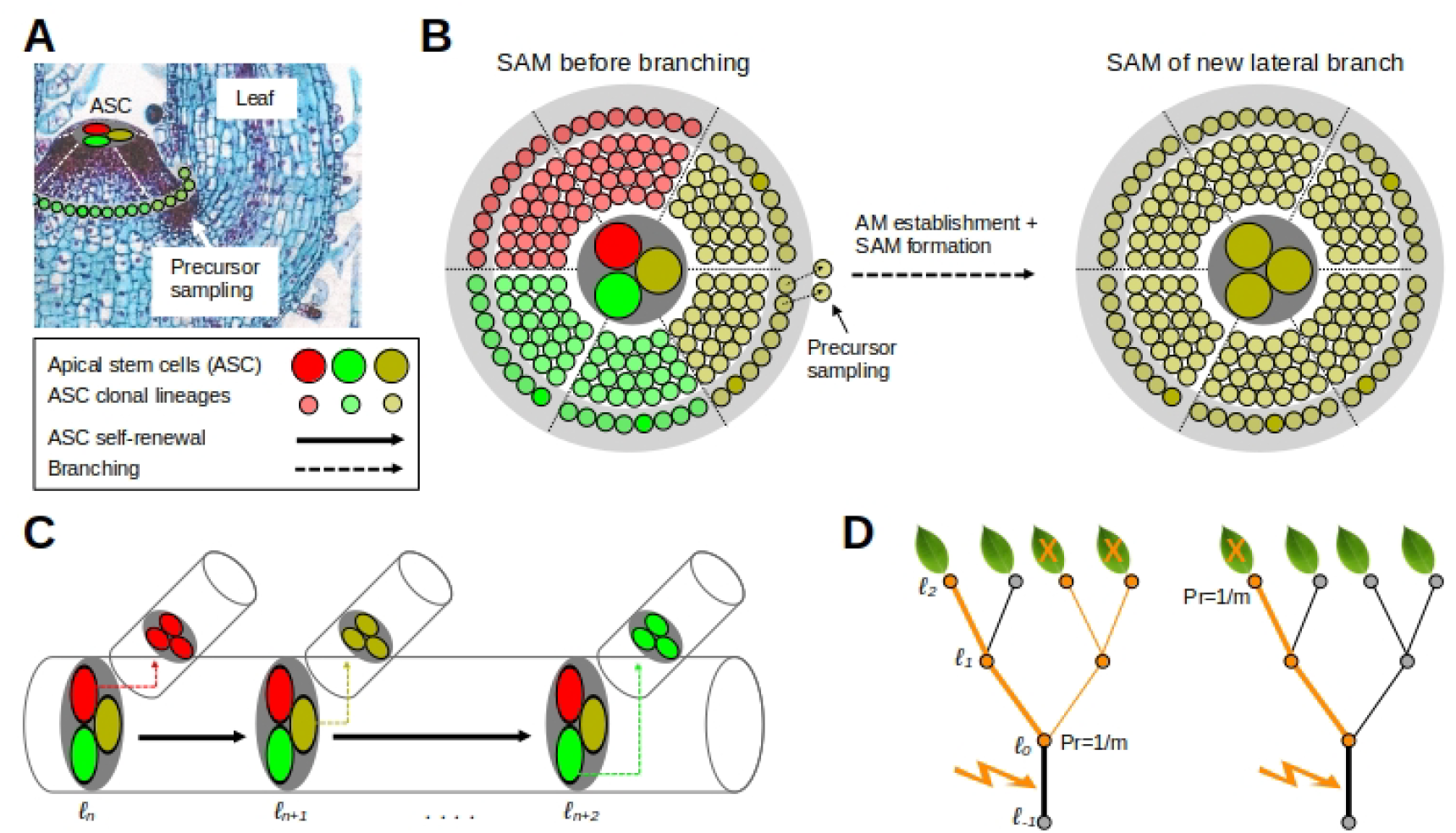
Branching model and ASC sampling. **A**. Side-view of the shoot apical meristem (SAM). The colored circles at the tip represent three (*m* = 3) apical stem cells (ASC). Clonal descendants of these ASCs are shown in corresponding colors at the SAM periphery. The arrow points to the site of axillary meristem (AM) establishment. **B**. A schematic top-view of the SAM. The tree clonal sectors are shown just before branching, with each sector corresponding to one of the ASC at the tip. During branching, precursors are “sampled” from the SAM periphery to acts as founders for the SAM of the new lateral branch or organ. The cellular composition of the new SAM reflects the proportional sampling of these sectors. Due to strong spatial constraints, precursors likely sample only one ASC via its clonal descendants, particularly if *m* is small. **C**. Under the cellular model shown in (B.) it is reasonable to assume a single ASC bottleneck model for shoot branching. The schematic shows how each branch is founded by a different ASC, although founder selection is probabilistic (see D.). **D**. Shown is the fate of a mutation arising in the bottom segment of the tree (see lighting bolt). How the mutations propagates to the crown tips depends on the sampling probabilities along the branching paths. Left panel: At branch point *ℓ* = 0 the mutation is selected into a lateral branch with probability 1*/m*, where it becomes fixed in all sub-trees descending from that branch point. At the same time, the mutant cell propagates further along the main stem, but in this case fails to be sampled at the terminal tip of the main step. Right panel: An alternative outcome, where the mutation fails to be selected into the lateral branch at *ℓ* = 0, but propagates all the way to the tip of the main stem. The sampling probabilities of a mutation arising on any segment *s* to be observed in the tips of the crown are derived in the main text.

Lateral branches arise from axillary meristems (AMs), which form in the axils of developing leaves from small groups of SAM peripheral cells (Shi et al. (2016)). Although multiple peripheral cells can initiate an AM, sampling is spatially restricted and often captures only a single ASC clonal sector per layer (Johannes (2025); Burian et al. (2016); He et al. (2024)) (Fig. 3B,C). We therefore adopt a *single-ASC bottleneck model* at each branching event: one ASC (per layer) is probabilistically chosen (via its daughter cells at the SAM periphery) to found the new meristem.

We acknowledge that this branching model is a simplification as precursor sampling can span clonal boundaries and momentarily incorporate multiple ASC lineages, leading to polyclonal AMs (e.g., Bossinger and Smyth (1996); Burian et al. (2016)). Indeed, Burian et al. (2016) estimate that roughly 39% of L1 AMs in Arabidopsis receive input from more than one ASC. However, subsequent branching events create additional bottlenecks that rapidly dilute mixed founder populations down to a single surviving lineage, as shown by Johannes (2025). Moreover, in Appendix A we demonstrate that allowing any small number of ASCs to found each new meristem leads to the same effective propagation behavior as our single-cell model. Thus, the single-ASC bottleneck captures the essential dynamics of lineage fixation and somatic drift while retaining analytic simplicity. We now proceed to develop the mutation model, starting with a single SAM layer before extending to all three.

#### ASC sampling probabilities during branching

Consider a mutation arising within the ASC population on segment *s* (Fig.3D). Above this segment lie *B*_*s*_ = *D* − *s* branch points where lateral shoots are formed. At each of these exactly one of the *m* ASC is randomly selected via its clonal descendants at the SAM periphery to act as a founder to a new lateral meristem (see Appendix A for a relaxation of this assumption). The continuing axis itself terminates in a final branch point where terminal organs such as fruit or leaves are formed (Fig.3D). Similar to lateral branching, we treat this terminal branch point as a cellular bottleneck, where one ASC founding cell is drawn uniformly from the *m* remaining cells. To be detected in a leaf or fruit, a mutation in *s* must therefore be selected in any one of the *B*_*s*_ lateral-branch draws or in the terminal founder draw for a total of *B*_*s*_ + 1 independent sampling events. Label these sampling events *j* = 1, 2, …, *B*_*s*_ + 1 and let

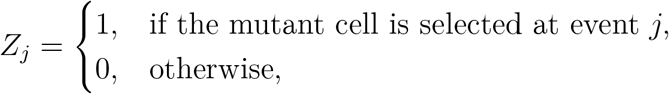

with each *Z*_*j*_ ∼ Bernoulli(1*/m*) (Fig.3D). Using this formalism, it is useful to define two key probabilities:

1. Lateral-branch entry probability: The chance the mutation enters a lateral branch at least once among the *B*_*s*_ branch points (ignoring the final draw) is

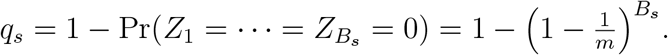
2. Unique-event detection probability: The chance the mutation is selected in *any* of the *B*_*s*_ + 1 draws (including the terminal founder) is

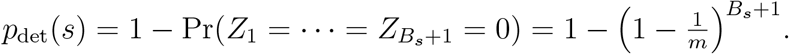

Since the probability of failing all *B*_*s*_ branch draws is 1 − *q*_*s*_, one can write

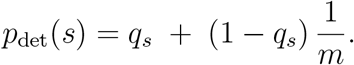

This decomposition partitions mutation detection into two disjoint events: “selection into a lateral branch” (probability *q*_*s*_) or “no lateral selection but success at the terminal draw” (probability (1− *q*_*s*_)*/m*). These two shorthand probabilities, *q*_*s*_ and *p*_det_(*s*), capture the entire sampling dynamics and serve as the building blocks for all crown-wide mutation-burden expectations described below.

### Expected somatic mutation burdens

#### Per-tip mutation burden

In any internode segment *s* each of the *m* cells mutates according to a Poisson process with rate 2*Gµ* over time *t*_*s*_ = *T w*_*s*_(*λ*), so that the total mutational supply from the ASC population of single shoot is *m* 2*Gµ t*_*s*_. Only one of those *m* cells is sampled at the terminal founder (prob. 1*/m*), so thinning yields (2*Gµt*_*s*_) per segment. Summing over all segments from root to tip gives the expected per-tip mutation burden:

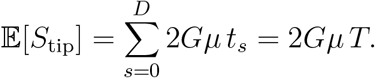

In practical terms, this would be the expected number of somatic mutations observed in a single sequenced lateral organ (e.g. a leaf or fruit), given the total age of the tree (*T*), the genome size (*G*) and the mutation rate (per-ASC per unit time).

#### Crown-wide mutation burden

Having derived the per-tip burden, it is of interest to find the expected somatic mutation burden in the tree crown as a whole. To do this, consider that each of the *C*_*s*_ constant-order axes contributes *m* stem cells mutating at rate 2*Gµ* over time *t*_*s*_ = *Tw*_*s*_(*λ*). Hence, the total mutation supply from all segments with index *s* is

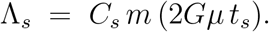

Recall that a mutation arising on a segment can escape laterally with probability

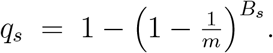

When this happens, it fixes in all 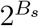 descendant tips of the lateral subtree (Fig.3D, left panel). Alternatively, the mutation fails to escape any lateral branch (probability 1 − *q*_*s*_) and must survive the final tip-founder draw with probability 1*/m*, thereby affecting exactly one tip (Fig.3D, right panel). The expected number of mutations arising on segment *s* that are inherited to the tips of the crown can therefore be expressed as

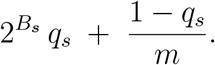

Summing over all segments and applying Poisson thinning gives

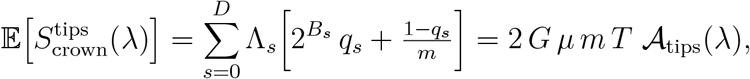

where the architectural component for total tip occurrences is

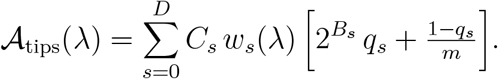

### Crown-wide unique-mutation burden

To count each mutation event observed in the crown tips at most once, regardless of how many tips inherit it, we collapse the subtree-amplification factor 2^*B*^*s* into a single “seen-or-not” outcome across the *B*_*s*_ lateral draws plus the terminal draw. Recall the unique-event detection probability of a mutation occuring o segment *s*

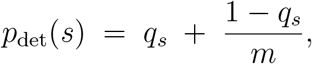

then the expection becomes

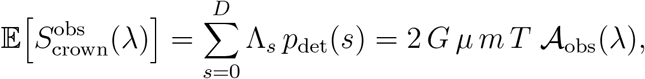

with the architectural component being defined by

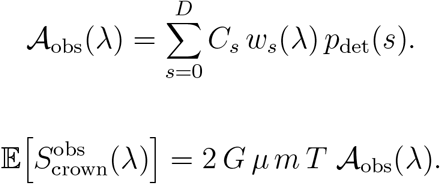

### Remark on polyclonal founding

Although we have built the above with a single-ASC bottleneck for clarity, empirical studies have found that axillary meristems are occasionally founded by multiple stem cells (e.g. Burian et al. (2016); Bossinger and Smyth (1996); Johannes (2025)). However, one can show (Appendix A) that sampling any fixed number *k* ≪ *m* of ASCs at each branch point leads to the same *net* propagation probability 1*/m* per branch–and–fixation cycle. Thus all of our tip- and crown-wide burden formulas - and their architectural dependence via A_obs_(*λ*) - remain exactly as stated.

### Simulation of crown-wide mutation burden

To validate the derived expectations, we implemented a Monte Carlo simulator that, for each combination of *D, m*, and *λ*, performs three steps: **1**. allocates segment times via the tilt function. **2**. draws Poisson-distributed mutational inputs in each segment for both tip-specific and crown-wide processes, and **3**. propagates each mutation through the *B*_*s*_ lateral-branch and terminal founder bottlenecks exactly as in the model. We find that the simulation closely agree with the theoretical expectations (Fig. 4, Table S1).

**Figure 4:**
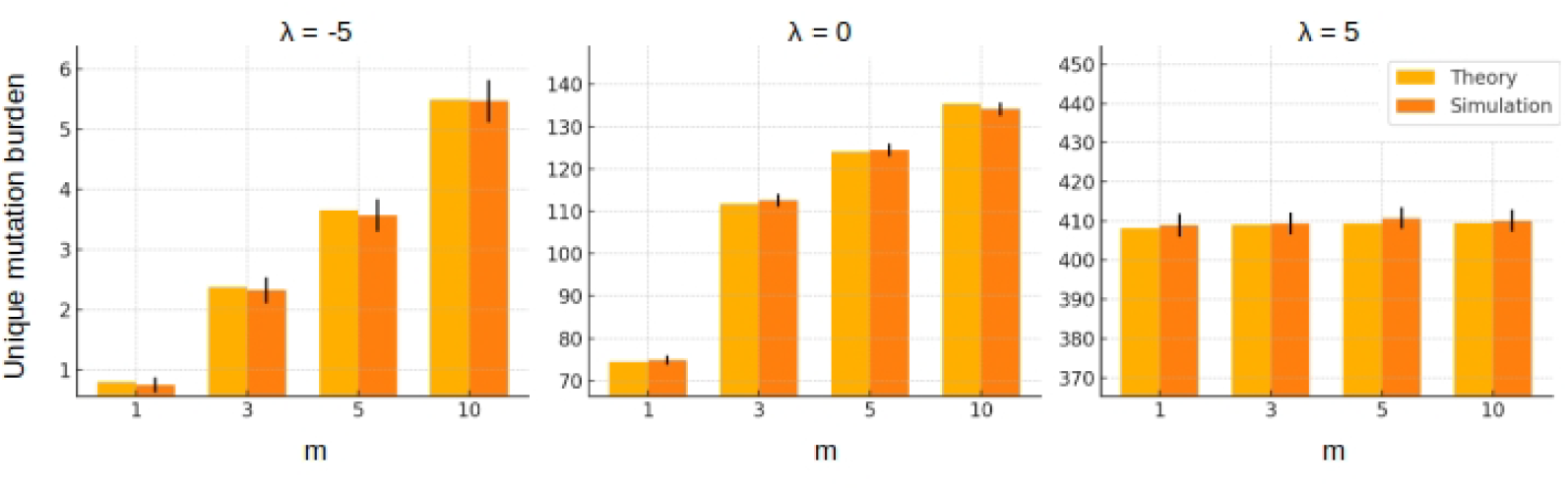
Simulation study. Evidence that the theoretical expectations agree with stochastic simulations. Example result from simulations of the crown-wide unique-mutation burden, 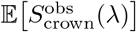, for *D* = 10, *m* ∈ {1, 3, 5, 10} and *λ* ∈ {−5, 0, 5}. Error bars are empirical 95% confidence intervals from 100 replicate runs. Full simulation results for a wider range of values can be found in Table S1.

### Architectural amplification of somatic mutation burden

The above results highlight that somatic mutation burden in a tree crown is partly shaped by branching architecture. Because the prefactor 2 *G µ m T* in the expectations is constant for a given tree at the time of sampling, the expected somatic mutation burden is simply proportional to the architectural components:

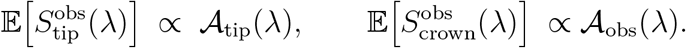

To quantify the impact of branching architecture, via _obs_, we examine how extreme and intermediate time-allocation regimes suppress or amplify mutational load.

#### Fold-change in burden between architectural extremes

Recall the sticktree and the star-tree architectural extremes. When *λ* → −∞ (stick-tree), all time is devoted to the basal segment *s* = 0, so that the architectual component becomes

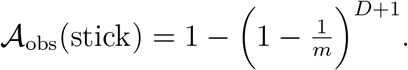

Conversely, as *λ* → + ∞ (star-tree), all time goes to the terminal segment *s* = *D*, yielding

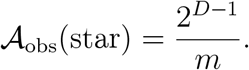

Because *𝒜*_obs_(*λ*) is a convex combination of the segment-wise terms

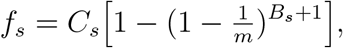

it always satisfies

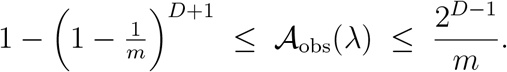

To appreciate the magnitude of differences in mutational burden that can be expected when comparing the two architectural extremes, we calculate their ratio

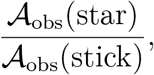

as a function of *m* and *D*. As shown in Fig.5A, star-trees can amplify somatic mutation burden in the tree crown by several orders of magnitude relative to stick-trees, even if the tree age *T* and the number of terminal tips 2^*D*^ are identical.

#### Fold-change in burden relative to uniform allocation

An alternative to evaluating the impact of branching architecture is to assess how departures from uniform growth suppress or amplify the crown-wide burden. When *λ* = 0, time is spread uniformly over all internode segments (*w*_*s*_ = 1*/*(*D* + 1)), yielding a baseline *𝒜*_obs_(0). Let us define

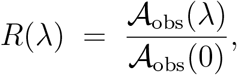

so that *R*(0) = 1, *R <* 1 indicates suppression, and *R >* 1 indicates amplification relative to the uniform case. Figure 5(B and C) illustrate that even modest tilts (| *λ* | ≈1) away from uniform growth can change the expected burden by a factor of 2–5, depending on *D* and *m*.

**Figure 5:**
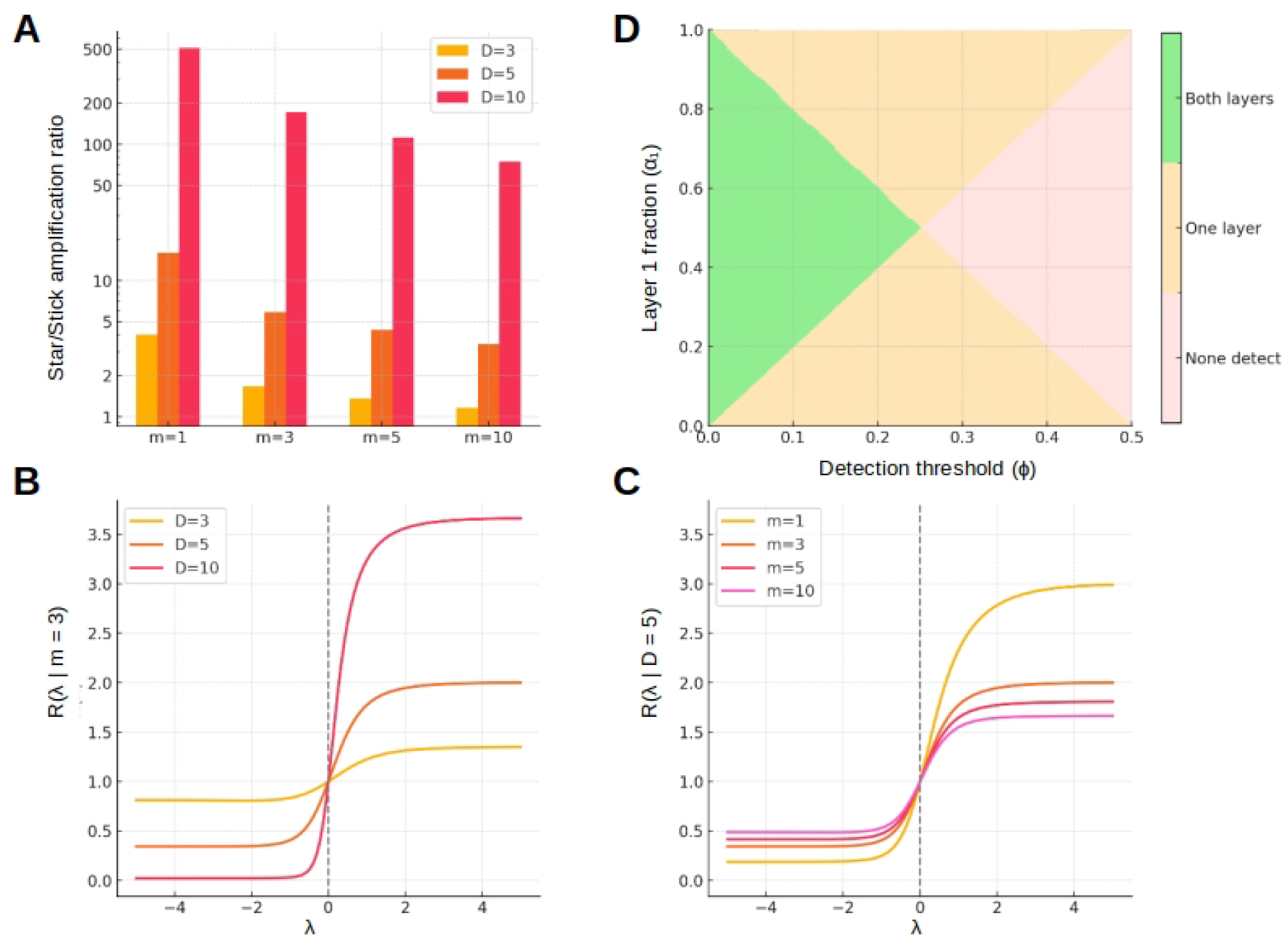
Architectural impact on somatic mutation burden. **A**. Fold-change differences in the unique-mutation burden between the star-tree architecture relative to the stick-tree archiecture plotted on a *log* scale. The magnitude of the fold-change depends mainly on the depth *D* of the tree, but also on the ASC population size. **B**. For a fixed ASC (*m* = 3), fold-change difference in the unique-mutation burden for different values of *λ* to a baseline architecture with uniform growth (*λ* = 0). **C**. For a fixed depth (*D* = 5), fold-change difference in the unique-mutation burden for different values of *λ* relative to a baseline architecture with uniform growth (*λ* = 0). **D**. Impact of variant allele threshold (*x*-axis) in sequencing data of lateral organs (e.g. fruits) on the ability to observe mutations originating in any of the three meristematic layers. For visual simplicity, we assume that only two layers contribute cells to the sequenced sample. Depending on the threshold, mutations can be detected either from both layers (green), one layer (beige) or no layer (pink).

### Impact of cell-layer structure and detection thresholds

So far we have ignored the layer-structure of the SAM. As already described above, the SAM is stratified into three histological layers, named L1, L2 and L3, each giving rise to different tissues (epidermis, subepidermal layers, pith, etc.)(Lyndon (1998)). When we sample a leaf or fruit in bulk, cells from these three layers mix in unknown proportions *α*_1_, *α*_2_ and *α*_3_ with

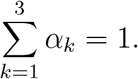

Because these layers are developmentally distinct, they may accrue somatic mutations at different per-base rates *µ*_*k*_ (Goel et al. (2024), even though they share the same branching architecture {*C*_*s*_, *B*_*s*_}, time allocation *w*_*s*_(*λ*), genome size *G* and presumably also the same ASC population size *m*. If we could sequence each layer separately, for example by laser-capture microdissection or layer-enriched protocols (e.g. Goel et al. (2024)), we would recover unique mutation lists from each layer and could estimate *µ*_*k*_ directly (Johannes (2025)):

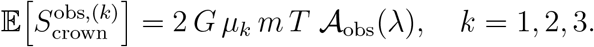

Summing gives the full crown burden across layers:

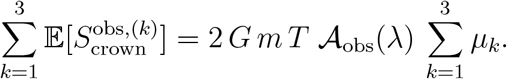

However, most empirical studies have relied on bulk sequencing of leaves or fruits, in which DNA from all three layers is mixed. Somatic mutations thus appear as variant alleles at frequency

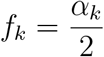

in diploid tissue. Standard variant callers typically require a minimum allele frequency *ϕ* (e.g. 5–10%) to distinguish true mutations from sequencing error. Hence, any mutation arising in layer *k* can be detected only if

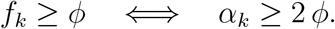

We capture this with the indicator

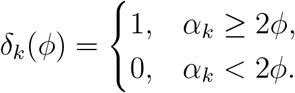

Under this bulk-sequencing constraint, layer *k* effectively contributes only its share of mutations if it exceeds the allele-frequency cutoff. The resulting expected unique-event burden in the bulk sample becomes

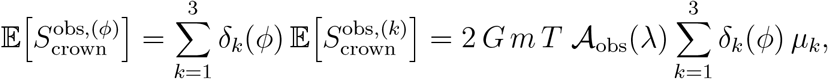

where the detection penalty on *µ* scales linearly in the expectation. Because 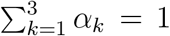, this mixture automatically weights each layer’s mutation rate by its cell–fraction contribution and excludes any layer whose mutations fall below the calling threshold. In practice, layers with small *α*_*k*_ (for example, L3 in many species) may become virtually invisible in bulk data (Fig.5D), biasing somatic-mutation estimates toward the rates of the major layers.

## Conclusion

Somatic mutations arise as a natural consequence of cell division during development, and their accumulation in long-lived plants has broad implications for ecology, evolution, and genome stability. Surprisingly, genomic studies have consistently reported low somatic mutation counts in mature trees (Johannes (2024)). Based on a rough calculation that did not take branching architecture into account, Schoen and Schultz (2019) argued that these counts should be several orders of magnitude higher, and attributed this discrepancy to errors in ascertaining mutations from sequencing data. While incomplete detection due to low allele frequencies in bulk sequencing may contribute, recent studies using layer-enriched or long-read calling approaches have yielded similar results (Goel et al. (2024); Johannes (2025); Xian et al. (2025)), suggesting that technical artifacts are not the primary explanation.

Instead, we propose that the spatio-temporal structure of the tree crown itself strongly suppress the accumulation of fixed somatic mutations. Because branching events are accompanied by tight cellular bottlenecks, and because terminal branches often share long portions of their developmental history, many cell divisions do not generate new, independently traceable mutations. Instead, the degree of developmental path-sharing among branches plays a key role in determining the likelihood that a mutation arising in a stem cell lineage becomes fixed and observable in the crown.

This same logic applies to somatic epimutations, which typically occur at much higher rates than DNA sequence mutations (Becker et al. (2011); Schmitz et al. (2011); van der Graaf et al. (2015); Hazarika et al. (2022); Yao et al. (2023)), but exhibit similar clonal inheritance in meristematic tissues (Hofmeister et al. (2020); Shahryary et al. (2020)). Although epimutations may revert over long timescales (van der Graaf et al. (2015)), such reversions are rare over the lifespan of a tree. Thus, our modeling framework extends naturally to both types of heritable molecular change, requiring no reparamterization.

Our model generates several empirically testable predictions. For example, trees with more path-sharing among terminal branches, those in which branching occurs late in development, are expected to exhibit lower mutation burdens than trees in which branches diverge early and grow independently. As a result, mutation accumulation should vary with branching strategy even when crown depth and number of tips are held constant. Moreover, because each branching event samples from a limited set of stem cell lineages, mutation accumulation should saturate with developmental depth. That is, adding more branching events beyond a certain point contributes little additional diversity, as the number of distinct ASC lineages that can be sampled is limited. This saturation effect implies that mutation counts should increase sublinearly with path length. Within a single tree, mutation sharing between terminal branches should reflect their developmental proximity: branches that diverged more recently should share more mutations than those that separated earlier. These predictions are amenable to testing through phylogenetic analysis of branches or direct quantification of mutation burden across trees with contrasting crown architectures. Furthermore, they suggest that experimentally altering branching patterns - for instance, through pruning or hormone treatments - should modulate mutation accumulation in predictable ways.

A caveat of our current model is that it focuses exclusively on mutations arising in the shoot apical meristem. This emphasis reflects the fact that SAM-derived mutations are more likely to become fixed due to the cellular bottlenecks at branching events, occur at high enough cellular frequencies to be reliably detected through genomic sequencing, are more amenable to mathematical modeling, and are likely to be functionally significant. By contrast, mutations that arise post-SAM, during organogenesis or later development, are expected to occupy small, localized sectors. Their distribution is less predictable and more difficult to model in relation to overall tree architecture. Neverthless, because such mutations do occur, our model offers a lower bound on the cumulative burden of somatic mutations in trees.

Taken together, our theoretical insights emphasize that tree architecture is not merely a consequence of growth, but also a developmental scaffold that governs the accumulation and distribution of somatic (epi)mutations. By linking growth form with molecular evolution, this framework opens new directions for studying how life-history strategies shape genetic and epigenetic diversity in long-lived plants.

## Appendix Robustness of somatic burden to multi-ASC sampling

In the main text, we assumed a single-ASC bottleneck at each branching event. Here we generalize to sampling *k* stem cells (with 1 ≤ *k* ≪ *m*) at each bottleneck and show that the architectural dependence of somatic mutation burden remains unchanged, even accounting for the terminal-segment fixation nuance.

## Notation

- *m*: total number of apical stem cells (ASCs) per SAM.
- *k*: number of ASCs drawn without replacement at each lateral branch point or terminal event.
- *D*: crown depth; *s* = 0, 1, …, *D*.
- *B*_*s*_ = *D* − *s*: number of lateral branches above segment *s*.
- *T, G, µ, C*_*s*_, *w*_*s*_(*λ*): as defined in the main text.

## Seeding probability for drawing *k* ASCs

A mutation present in exactly one of the *m* ASCs on segment *s* enters the newly established SAM if at least one of the *k* drawn founders is the mutant:

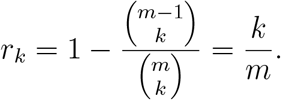

Immediately after seeding, there are *k* coexisting founder lineages, one of which carries the mutation, each at frequency 1*/k*.

## Branch-and-fix cycle for non-terminal segments

For segments *s* < *D*, each lateral branch point entails a two-step process:

1. **Seeding draw** of *k* founders (probability *r*_*k*_ to include the mutant).
2. **Fixation draw** among those, *k* lineages will be selected at the next bottleneck (probability 1*/k* that the mutant lineage is chosen).

The net propagation probability per branch-and-fix event is therefore:

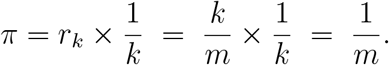

## Unique-detection probability for segments *s < D*

Above segment *s* there are *B*_*s*_ lateral branch events and one final tip event, totaling *B*_*s*_ + 1 independent branch-and-fix trials, each succeeding with probability 1*/m*. Thus,

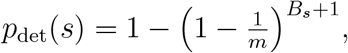

identical to the single-ASC case.

## Terminal-segment (*s* = *D*) nuance

Mutations arising on the terminal segment have no subsequent lateral branch to seed, only the final draw of *k* ASCs, which is both the seeding and fixation event. They propagate if at least one of the *k* founders is the mutant:

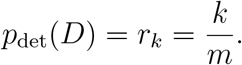

In the single-ASC model this was taken as 1*/m*. Replacing only this term adjusts the architectural sum by a factor *k* for the *s* = *D* contribution:

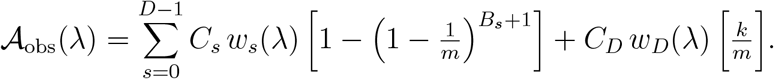

Because *k* ≪ *m* and under most architectures *w*_*D*_(*λ*) remains small, this correction is negligible, preserving all qualitative conclusions.

## Expected crown-wide unique mutation burden

The total mutational supply on segment *s* remains:

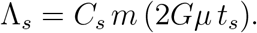

Thus:

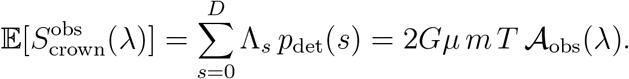

## Conclusion

Sampling *k* ASCs (with *k* ≪ *m*) per bottleneck yields the same net propagation probability 1*/m* for all non-terminal segments. The terminal-segment correction of *k/m* is negligible under typical architectures, leaving the architectural amplification factor A_obs_(*λ*) and all downstream conclusions unchanged.

## Acknowledgments

I thank Lutz Johannes and Tomàs Johannes-Colomé for their help in generating Fig. 1A. Thanks also to Benjamin Werner for his quick feedback. This work was partly funded by the Bundesministerium für Bildung und Forschung (Project: epiSOMA).

